# Clustering-mediated activation of Cdc42 GTPase antagonized by GAPs in fission yeast

**DOI:** 10.1101/2020.06.02.130336

**Authors:** Iker Lamas, Nathalie Weber, Sophie G Martin

## Abstract

The small GTPase Cdc42 is critical for cell polarization in eukaryotic cells. In rod-shaped fission yeast *Schizosaccharomyces pombe* cells, active GTP-bound Cdc42 promotes polarized growth at cell poles, while inactive Cdc42-GDP localizes ubiquitously also along cell sides. Zones of Cdc42 activity are maintained by positive feedback amplification involving the formation of a complex between Cdc42-GTP, the scaffold Scd2 and the guanine nucleotide exchange factor (GEF) Scd1, which promotes the activation of more Cdc42. Here, we use the CRY2-CIB1 optogenetic system to recruit and cluster a cytosolic Cdc42 allele at the plasma membrane and show that this leads to its moderate activation also on cell sides. Surprisingly, activation of CRY2-Cdc42 does not individually depend on Scd1 or the GEF Gef1. We show that activated Cdc42 clusters at cell sides are able to recruit Scd1, dependent on the scaffold Scd2. However, Cdc42 activity is not amplified by positive feedback and does not lead to morphogenetic changes, due to antagonistic activity of the GTPase activating protein Rga4 on cell sides. Thus, the cell architecture is robust to moderate activation of Cdc42 at cell sides.

## Introduction

In eukaryotes, the small Rho-family GTPase Cdc42 is a highly conserved regulator of cell morphogenesis, proliferation and differentiation. Prenylation of Cdc42’s C-terminal CAAX motif underlies its association with the plasma membrane, where it functions as a molecular switch that alternates between GTP-bound, active and GDP-bound, inactive states. Activation of Rho GTPases relies on the activity of guanine nucleotide exchange factors (GEFs), while their intrinsic GTPase activity is enhanced by GTPase activating proteins (GAPs) to return them to the inactive state. GDP-bound Cdc42 also binds guanine-nucleotide dissociation inhibitors (GDI), which both block the exchange of GDP by GTP and solubilize Cdc42-GDP in the cytosol.

In the fission yeast *Schizosaccharomyces pombe,* Cdc42 is active at sites of polarized growth during vegetative and sexual life cycles. GTP-loading is promoted by two GEFs, Scd1 and Gef1. Scd1, which localizes to cell poles, receives information from the upstream Ras1 GTPase signal and mediates feedback control through the scaffolding activity of Scd2 (Bendezu et al., 2015; Kelly and Nurse, 2011; Lamas et al., 2020). For this, Scd1 forms a quaternary complex with Cdc42-GTP, the Pak1 kinase effector and Scd2 (Chang et al., 1994; Endo et al., 2003), which leads *in vivo* to the positive feedback activation of other Cdc42 molecules, as shown in our recent work using optogenetic strategies (Lamas et al., 2020). The second GEF, Gef1, which localizes to cell poles only in some conditions, promotes Cdc42 activation in response to stress and becomes essential only in absence of Scd1 (Coll et al., 2003; Hirota et al., 2003; Tay et al., 2018; Vjestica et al., 2013). Three GTPase activating proteins (GAPs) Rga4, Rga6 and Rga3 enhance the intrinsic GTP hydrolytic activity of Cdc42 (Das et al., 2007; Gallo Castro and Martin, 2018; Revilla-Guarinos et al., 2016). Rga4 and Rga6 GAPs localize at cell sides, where growth does not occur in non-stressed cells, whereas Rga3 localizes at sites of active growth (cell poles). Fission yeast cells also express a GDI, called Rdi1, though Cdc42 localization and dynamics are not strongly perturbed in its absence (Bendezu et al., 2015; Nakano et al., 2003).

Recently, optogenetic studies revealed a novel mechanism that triggers the activation of small GTPases in mammalian cells: Human Rac1 and RhoA, which belong to the same Rho GTPase family as Cdc42, were shown to become active at the cell cortex upon light-dependent cytosolic clustering (Bugaj et al., 2013). In these experiments, the small GTPases were fused to CRY2PHR, the photolyase homology region of *A. thaliana* cryptochrome 2, which oligomerizes upon blue light exposure. Artificially clustered RhoA induced RhoA signalling-dependent cytoskeletal re-organization and membrane retraction in human cells, suggesting that oligomerization promotes RhoA activation (Bugaj et al., 2013). Ras and Ras-like GTPases are well known to form nanoclusters and dimers at the membrane to activate signal transduction (Inouye et al., 2000; Kang et al., 2010; Nussinov et al., 2020). Several Rho-family GTPases, including RhoA, Rac1, Rac2 and Cdc42, were also shown to form dimers or oligomers through homophilic interactions of their polybasic region adjacent to the C-terminal CAAX motif (Zhang et al., 2001; Zhang and Zheng, 1998). While oligomerization of GTP-bound Cdc42 and Rac1 increases their GTPase activity *in vitro,* the physiological relevance of clustering of these small GTPases remains to be investigated (Zhang and Zheng, 1998). *In vivo*, Rac1-GTP oligomers have been shown to contain several dozen Rac1 molecules together with charged phospholipids and appear to promote signal transduction (Maxwell et al., 2018; Remorino et al., 2017; Zhang et al., 2001). Cdc42 also forms nanoclusters in *Saccharomyces cerevisiae* cells (Slaughter et al. 2013; Sartorel et al. 2018). These nanoclusters show an anisotropic distribution: they accumulate and exhibit larger sizes at cortical sites of polarized growth, in a manner dependent on the scaffold protein Bem1 and anionic membrane lipids (Meca et al., 2019; Sartorel et al., 2018). Because Bem1 also acts as scaffold that bridges Cdc42-GTP to its GEF, thereby promoting positive feedback activation of Cdc42, Cdc42 nanoclusters may promote feedback Cdc42 activation, though this has not been tested.

In this work, we used an artificial optogenetic strategy to induce the recruitment and clustering of Cdc42 at the plasma membrane of fission yeast cells. We built on our recent work that used the CRY2-CIB1 optogenetic system to probe the positive feedback of Cdc42 (Lamas et al., 2020). The CRY2-CIB1 system relies on the blue light-induced binding of CRY2PHR (simply denoted CRY2 below) to the N-terminal part of CRY2-binding partner CIB1 (CIBN) (Kennedy et al., 2010). Blue light also induces the formation of CRY2 oligomers (Bugaj et al., 2013). We fused CRY2 to a cytosolic allele of Cdc42 (Cdc42^ΔCaaX^) and co-expressed CIBN linked to the membrane-associated RitC anchor. Cortical recruitment of a GTP-locked, constitutively active Cdc42 allele (CRY2-Cdc42^Q61L,ΔCaaX^) led to the Scd2-dependent co-recruitment of its GEF Scd1 and accumulation of endogenous Cdc42, demonstrating feedback amplification. Surprisingly, cortical recruitment of CRY2-Cdc42^ΔCaaX^ (not GTP-locked) also induced the co-recruitment of Scd2 (Lamas et al, 2020), suggesting activation of CRY2-Cdc42^ΔCaaX^. Here, we confirm that CRY2-dependent clustering promotes Cdc42 activation at lateral sites, where Cdc42 is normally inactive. Activated clustered Cdc42 is able to recruit its GEF through the scaffold Scd2, suggesting that positive feedback is initiated. However, the activation is efficiently countered by Rga4 GAP-mediated Cdc42 inactivation, and does not lead to cell shape alteration, showing the robustness of the cell polarization system.

## Results and discussion

### Weak activation of CRY2-Cdc42 at the cell cortex

To better characterize CRY2-Cdc42^ΔCaaX^, we first measured its kinetics of recruitment to CIBN-RitC at the plasma membrane. Similar to rates measured for CRY2, CRY2-Cdc42^ΔCaaX^ showed a half-time of protein recruitment to the cortex < 1 s and independent of the length of the blue light (488nm) pulses (30 GFP pulses of 50 ms = 0.92 s ± 0.24 s; 22 GFP pulses of 250 ms = 0.98 s ± 0.25 s; 17 GFP pulses of 500 ms = 0.99 s ± 0.31 s; Fig S1A-B). CRY2-Cdc42^ΔCaaX^ cells did not exhibit any morphological defects and grew in a bipolar fashion in the dark (Fig S1C). In blue-light, CRY2-Cdc42^ΔCaaX^ cells maintained their characteristic rod-shaped and continued growing from the cell tips (Fig S1D – green cells), while cells with activated CRY2-Cdc42^Q61L,ΔCaaX^ rounded up indicating isotropic growth (Fig S1D – blue cells; (Lamas et al., 2020)). These evidences initially suggested that the recruitment of CRY2-Cdc42^ΔCaaX^ to the cell cortex was innocuous and unable to bias the endogenous Cdc42 and its regulatory network.

We monitored the distribution of Cdc42-GTP using three GFP-tagged markers that specifically associate with Cdc42-GTP: the scaffold Scd2 (Endo et al., 2003; Wheatley and Rittinger, 2005), the CRIB bioreporter (Tatebe et al., 2008), and the Cdc42 effector Pak1, which also contains a CRIB domain. These proteins and probe are normally only detected at the poles and division sites of yeast cells, as well as weakly in the nucleus for the first two. As previously described (Lamas et al 2020), upon blue light-dependent recruitment of CRY2-Cdc42^ΔCaaX^ to the plasma membrane, Scd2-GFP formed stable foci at the cell sides, which increased progressively in intensity, while Scd2-GFP intensity decreased at cell poles (Fig 1A-B). CRIB-3GFP also formed dim, dynamic foci at the cell sides, which became visible 40-60s after light stimulation (Fig 1A-B). The CRIB-3GFP side signal was however weaker and delayed relative to that observed upon light-induced recruitment of GTP-locked CRY2-Cdc42^Q61L,ΔCaaX^ (Fig 1C; (Lamas et al., 2020)). The Pak1-sfGFP trace on cell sides also showed an upward trajectory, but was not statistically different from negative control after 90s, suggesting delayed recruitment below the detection threshold (Fig 1A-C). Together, these data suggest that the heterologous allele of Cdc42 within the CRY2-Cdc42^ΔCaaX^ system is transiently activated when recruited in clusters at the cell sides.

**Figure 1.**
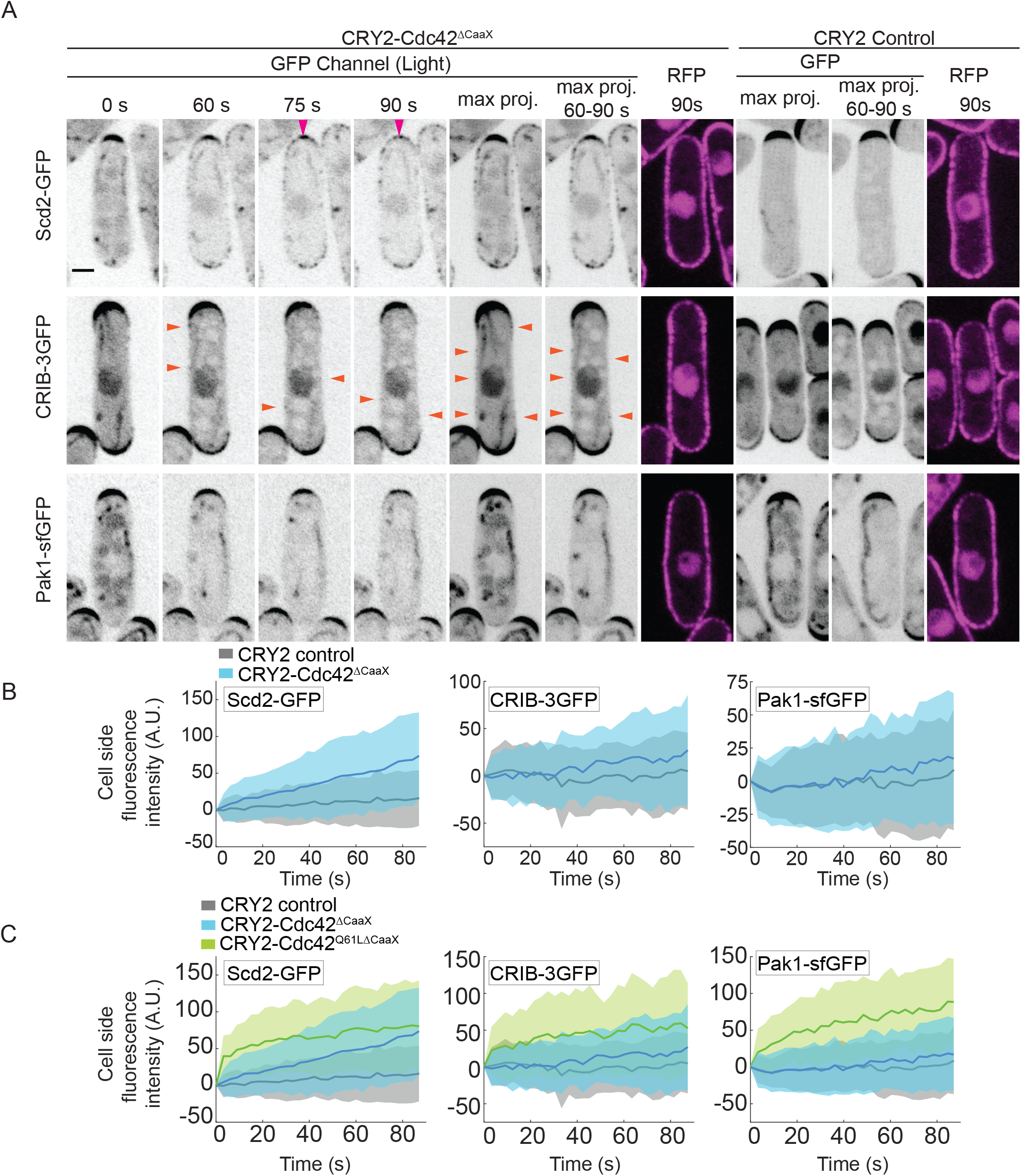
Ectopic sites of Cdc42 activation upon CRY2-Cdc42^ΔCaaX^ cell-sides recruitment. (A) Localization of Scd2-GFP, CRIB-3GFP and Pak1-sfGFP in CRY2-Cdc42^ΔCaaX^-expressing cells (B/W inverted images). The GFP max projection (“max proj.”) images show GFP maximum-intensity projections of 30 time points over 90 s and illustrate best the side recruitment of GFP-tagged probes. The GFP max projection 60 – 90 s (“max proj. 60 – 90 s”) images show GFP maximum-intensity projections of 10 last time points over 30 s. Orange arrowheads point to lateral CRIB-3GFP signal. Pink arrowheads point to Scd2-GFP tip signal decrease. RFP images show the cortical recruitment of CRY2-Cdc42^ΔCaaX^ and CRY2 at the end of the time lapse. The internal globular and filamentous signal is due to cellular autofluorescence. (B) Quantification of GFP signal intensities of Scd2-GFP, CRIB-3GFP and Pak1-sfGFP at cell sides in WT cells. *N* = 3; *n* > 20 cells per experiment; *p*^Scd2-GFP^ = 1.55e^−8^; *p*^CRIB-3GFP^ = 0.019; *p*^Pak1-sfGFP^ = 0.168. (C) Comparison of data from panel (B) with CRY2-Cdc42^Q61L,ΔCaaX^ -induced cell-side re-localization of Scd2-GFP, CRIB-3GFP and Pak1-sfGFP (data from (Lamas et al., 2020)). In all graphs, thick line = average; shaded area = standard deviation; WT, wild type; A.U., arbitrary units. Bars = 2 μm. Associated trace analysis is shown in Fig S3A.

As an alternative strategy to increase Cdc42 levels at the plasma membrane, we overexpressed Cdc42. In this experiment, we used the functional, internally tagged Cdc42-mCherry^SW^ allele (Bendezu et al., 2015) expressed under the *p*^*act1*^ promoter in cells lacking the endogenous gene, which allowed us to quantify the global increase in expression levels at 3.3-fold (Fig S2A-B). The Cdc42 level increase was roughly uniform around the cell cortex (not shown). Cdc42 overexpression also led to a 1.2-fold increase in expression of the CRIB-3GFP reporter (expressed under the *p*^*pak1*^ promoter, Fig S2B). Cdc42 overexpression led to a small increase in CRIB signal at cell poles (even after correction by the 1.2-fold increase in probe expression) and a small increase in cell length (*p^act1^-cdc42* cell length = 14.2 ± 1 μm vs WT cell length = 13.4 ± 1.1 μm, t-test p-value = 2.8e-5; p^act1^-cdc42 cell width = 3.8 ± 0.3 μm vs WT cell width = 3.8 ± 0.3 μm, t-test p-value = 0.96), suggesting increased Cdc42 activity at cell poles. However, Cdc42 overexpression had no effect on CRIB-3GFP levels at cell sides (Fig S2C-D). We conclude that activation of CRY2-Cdc42^ΔCaaX^ on cell sides is not simply a consequence of overexpression but may be due to forced CRY2-dependent clustering of Cdc42.

### CRY2-Cdc42 activation still occurs in absence of either Cdc42 GEF

To probe the mode of CRY2-Cdc42^ΔCaaX^ activation, we repeated the optogenetic experiments above in strains lacking the Cdc42 GEF Scd1. In *scd1Δ* cells, Pak1-sfGFP was not detected at cell sides, as expected. We also observed only rare CRIB-3GFP dots, and no significant increase in CRIB levels at the sides of *scd1Δ* cells (Fig 2A-B), suggesting Scd1 may participate in CRY2-Cdc42^ΔCaaX^ activation. However, Scd2-GFP was still recruited to cell sides (Fig 2A-B), indicating Scd1 is not the sole activator of Cdc42 in these cells. We thus probed the role of the second Cdc42 GEF Gef1. In *gef1Δ* cells, Scd2-GFP accumulation at cell sides was not affected and exhibited similar re-localization dynamics as in WT cells (Fig 2C-D). We conclude that neither Cdc42 GEF is individually required for initial activation of clustered Cdc42 at the plasma membrane. It is possible that the two GEFs work redundantly in this situation. The lethality of *scd1Δ gef1Δ* double mutants unfortunately does not allow testing of this hypothesis (Coll et al., 2003; Hirota et al., 2003). Alternatively, Cdc42 clustering through CRY2 binding may promote its activation in GEF-independent manner.

**Figure 2.**
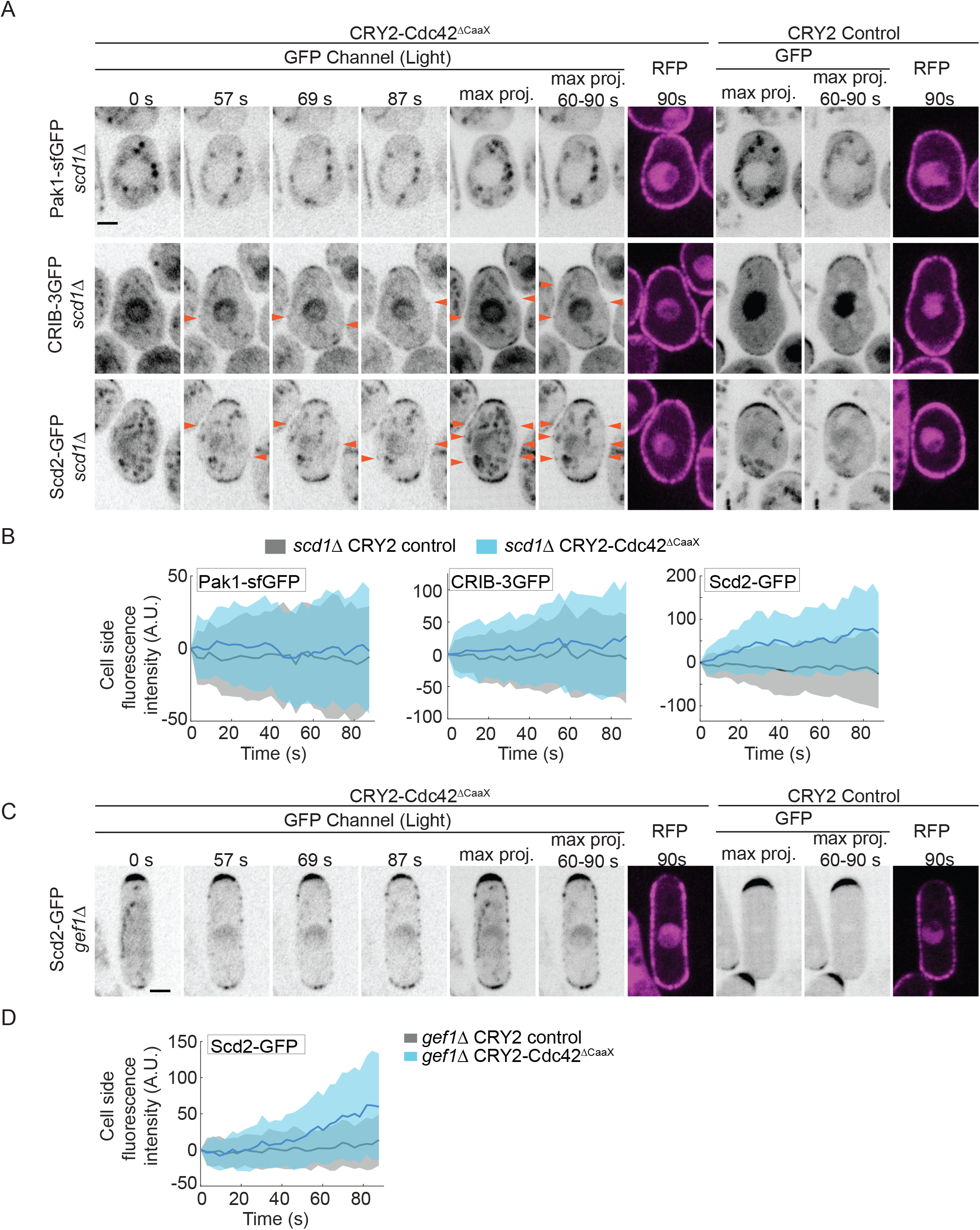
Role of Cdc42 GEFs in activation of CRY2-Cdc42^ΔCaaX^. (A) Localization of Scd2-GFP, CRIB-3GFP and Pak1-sfGFP in CRY2-Cdc42^ΔCaaX^-expressing *scd1Δ* cells (B/W inverted images). The GFP max projection (“max proj.”) images show GFP maximum-intensity projections of 30 time points over 90 s and illustrate best the side recruitment of GFP-tagged probes. The GFP max projection 60 – 90 s (“max proj. 60 – 90 s”) images show GFP maximum-intensity projections of 10 last time points over 30 s. Orange arrowheads point to lateral CRIB-3GFP and Scd2-GFP signal. RFP images show the cortical recruitment of CRY2-Cdc42^ΔCaaX^ and CRY2 at the end of the time lapse. The internal globular and filamentous signal is due to cellular autofluorescence. (B) Quantification of GFP signal intensities of CRIB-3GFP, Pak1-sfGFP and Scd2-GFP at cell sides of *scd1Δ* mutants. *N* = 3; *n* > 20 cells per experiment; *p*^Pak1-sfGFP^ = 0.172; *p*^CRIB-3GFP^ = 0.119; *p*^Scd2-GFP^ = 2.49e^−9^. (C) Localization of Scd2-GFP in CRY2-Cdc42^ΔCaaX^-expressing *gef1Δ* cells (B/W inverted images). Arrangement of images as in panel A. (D) Quantification of GFP signal intensity of Scd2-GFP at cell sides of *gef1Δ* mutants. *N* = 3; *n* > 20 cells per experiment; *p*^Scd2-GFP^ = 2.82e^−4^. In all graphs, thick line = average; shaded area = standard deviation. Bars = 2 μm. A.U., arbitrary units; WT, wild type. Associated trace analysis is shown in Fig S3B-C.

### CRY2-Cdc42 promotes recruitment of its GEF Scd1 in Scd2 scaffold-dependent manner

Because Cdc42-GTP promotes the recruitment of its GEF Scd1 for feedback amplification of Cdc42 activation (Lamas et al 2020), we probed whether CRY2-Cdc42^ΔCaaX^ induces Scd1 recruitment. Indeed, Scd1 formed weak, dynamic foci at cell sides upon blue-light activation, similar to the CRIB-3GFP foci described above (Fig 3A-B). The appearance of Scd1 dynamic foci at cell sides was also mirrored by a decrease of Scd1-3GFP at the cell tips (Fig 3C). Scd1 recruitment was dependent on the scaffold Scd2, as no cell side accumulation of Scd1-3GFP, nor decrease at cell tips, was detected in *scd2Δ* cells (Fig 3D-F). These data suggest that the activated CRY2-Cdc42^ΔCaaX^ is poised to trigger the positive feedback leading to recruitment of its GEF Scd1.

**Figure 3.**
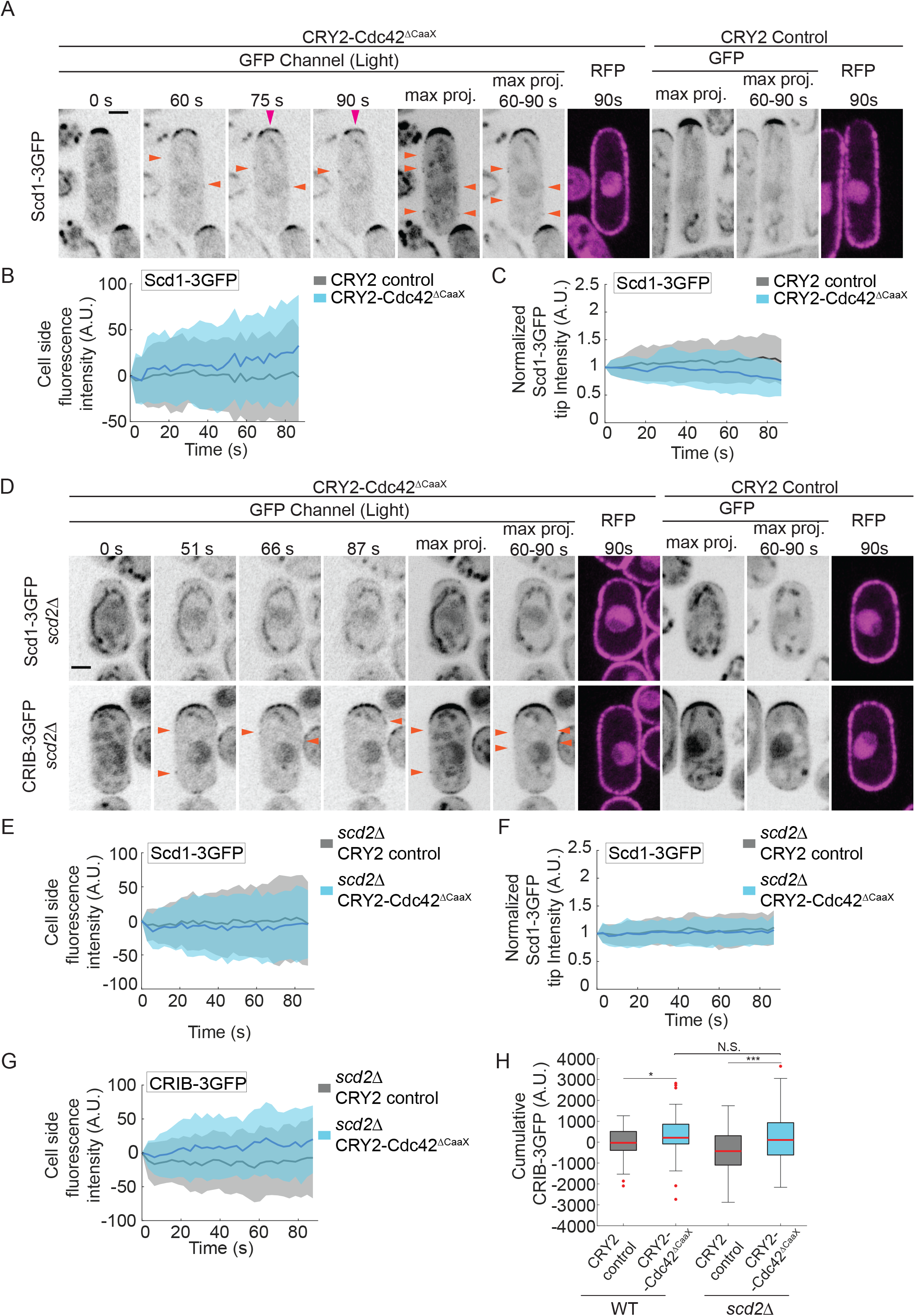
CRY2-Cdc42^ΔCaaX^ recruits the GEF Scd1 dependent on the scaffold Scd2. (A) Localization of Scd1-3GFP in CRY2-Cdc42^ΔCaaX^-expressing cells (B/W inverted images). The GFP max projection (“max proj.”) images show GFP maximum-intensity projections of 30 time points over 90 s and illustrate best the side recruitment of GFP-tagged probes. The GFP max projection 60 – 90 s (“max proj. 60 – 90 s”) images show GFP maximum-intensity projections of 10 last time points over 30 s. Orange arrowheads point to lateral Scd1-3GFP signal. Pink arrowheads point to Scd1-3GFP tip signal decrease. RFP images show the cortical recruitment of CRY2-Cdc42^ΔCaaX^ and CRY2 at the end of the time lapse. The internal globular and filamentous signal is due to cellular autofluorescence. (B) Quantification of GFP signal intensity of Scd1-3GFP at cell sides of WT cells. N = 3; n > 20 cells per experiment; p^Scd1-3GFP^ = 0.012. (C) Quantification of GFP signal intensity of Scd1-3GFP at cell poles of WT cells expressing CRY2-Cdc42^ΔCaaX^ or CRY2 (p = 2.14e-4). N = 3; n > 15 cells per experiment. (D) Localization of Scd1-3GFP and CRIB-3GFP in CRY2-Cdc42^ΔCaaX^-expressing *scd2Δ* cells (B/W inverted images). Arrangement of images as in panel A. Orange arrowheads point to lateral CRIB-3GFP signal. The internal globular and filamentous signal is due to cellular autofluorescence. (E) Quantification of Scd1-3GFP at cell sides of *scd2Δ* mutant. *N* = 3; *n* > 20 cells per experiment; *p*^Scd1-3GFP^ = 0.386. (F) Quantification of Scd1-3GFP at cell poles of *scd2Δ* mutants expressing CRY2-Cdc42^ΔCaaX^ or CRY2 (p = 0.37). N = 3; n > 15 cells. (G) Quantification of CRIB-3GFP at cell sides of *scd2Δ* mutants. *N* = 3; *n* > 20 cells per experiment; *p*^CRIB-3GFP^ = 7.67e^−4^. (H) Comparison of cumulative CRIB-3GFP intensities in WT and *scd2Δ* CRY2 and CRY2-Cdc42^Q61L,ΔCaaX^ cells; in *scd2*^*+*^ p^control vs CRY2-Cdc42^ = 0.019 (data from Figure 1B); in *scd2Δ* p^control vs CRY2-Cdc42^ = 7.67e^−4^ (data from panel G); p^CRY2-Cdc42 in WT vs *scd2Δ*^ = 0.4. In all graphs, thick line = average; shaded area = standard deviation. Bars = 2 μm. A.U., arbitrary units; WT, wild type. Associated trace analysis is shown in Fig S4.

Although *scd2* deletion abolished Scd1 recruitment, it did not substantially affect the accumulation of the CRIB probe (Fig 3D, 3G-H), indicating that CRY2-Cdc42^ΔCaaX^ is still activated in these cells. This observation is in agreement with the finding that CRY2-Cdc42^ΔCaaX^ is activated independently of Scd1. Because CRIB intensity on cell sides was not reduced in *scd2Δ* cells (Fig 3H), we conclude that the scaffold-dependent recruitment of the GEF by CRY2-Cdc42^ΔCaaX^ does not play a major role in amplifying Cdc42 activation at cell sides.

### The Cdc42 GAP Rga4 prevents isotropic growth of CRY2-Cdc42^ΔCaaX^ cells

If CRY2-Cdc42^ΔCaaX^ is activated at cell sides and recruits its own GEF, why is Cdc42 activity not further amplified by the positive feedback mechanism and does not lead to cell shape changes? Indeed, even long-term growth of CRY2-Cdc42 cells in the light did not change their cell length and width, or aspect ratio, similar to control CRY2 cells. By contrast, constitutive cortical localization of CRY2-Cdc42^Q61L,ΔCaaX^ by growth in light conditions led to significant increase in cell width and decrease in cell length, yielding a reduced aspect ratio (Fig 4A-C).

**Figure 4.**
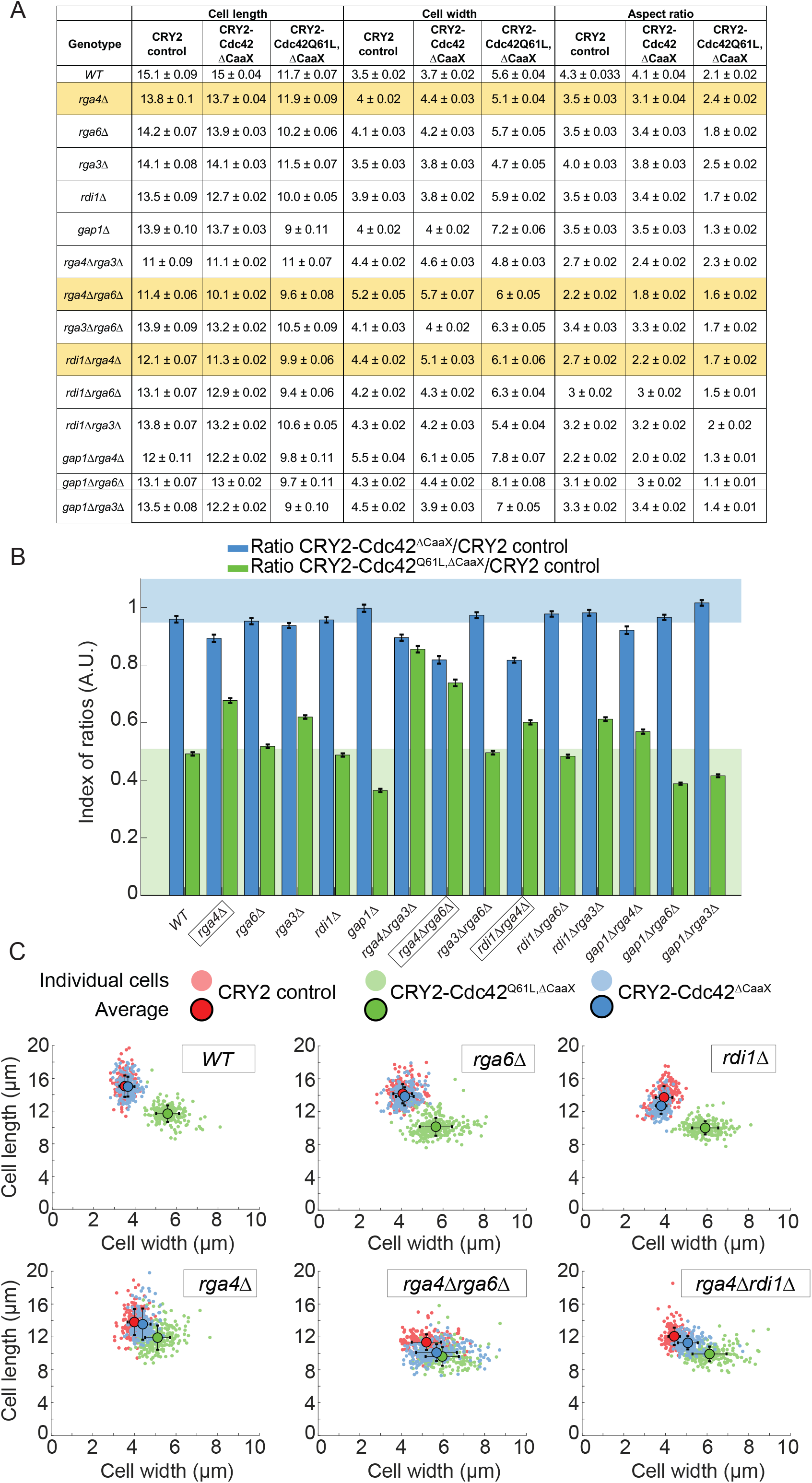
Rga4 GAP prevents growth on cell sides in CRY2-Cdc42^ΔCaaX^ cells. (A) Average cell length (μm), cell width (μm) and aspect ratio of CRY2, CRY2-Cdc42^ΔCaax^ and CRY2-Cdc42^Q61L,ΔCaaX^ mutants. For all mutant N = 2, n > 80 cells per experiment; except for *gap1Δ* mutants, N = 1, n > 80 cells per experiment. (B) Aspect ratios of CRY2-Cdc42^ΔCaax^ and CRY2-Cdc42^Q61L,ΔCaax^ -expressing cells grown in the light normalized to the aspect ratios of CRY2-expressing cells for all the tested mutants. Bars = standard error. The green background indicates expected reduction in aspect ratio upon CRY2-Cdc42^Q61L,ΔCaaX^ recruitment in WT cells. The blue background indicates absence of change in aspect ratio upon CRY2-Cdc42^Q61L^ recruitment. Note that all *rga4Δ* mutants fall in the white intermediate space. (C) Cluster plot of length and width in single cells of WT, *rga4Δ, rga6Δ, rga4Δrga6Δ, rdi1Δ and rga4Δrdi1Δ* mutants expressing CRY2, CRY2-Cdc42^ΔCaax^ or CRY2-Cdc42^Q61L,ΔCaaX^. Small dots = single cells; Large, dark dots = average; bars = standard deviation; A.U., arbitrary units; WT, wild type.

We hypothesized that the activation of CRY2-Cdc42^ΔCaaX^ at cell sides is rapidly counteracted by negative regulators. We focused our attention on the three Cdc42 GAPs Rga3, Rga4 and Rga6, the GDI protein Rdi1 and the Ras1 GAP Gap1. Rga3, Rga4 and Rga6 directly promote Cdc42-GTP hydrolysis (Das et al., 2007; Gallo Castro and Martin, 2018; Revilla-Guarinos et al., 2016). Rdi1 may promote Cdc42 extraction from the membrane, although it is largely dispensable for Cdc42 dynamics (Bendezu et al., 2015; Nakano et al., 2003). Gap1 directly promotes Ras1-GTP hydrolysis (Merlini et al., 2018). As Ras1 promotes Scd1 activation and is uniformly active at the plasma membrane in *gap1Δ* (Lamas et al., 2020; Merlini et al., 2016; Merlini et al., 2018), we hypothesized Scd1 activation on cell sides may be amplified in this mutant. We constructed single and most double deletion mutants expressing either of CRY2-Cdc42^ΔCaaX^, CRY2-Cdc42^Q61L,ΔCaaX,^ as positive control or CRY2 alone as negative control. We then measured the cell length and cell width of calcofluor-stained dividing cells after at least 14h of exponential growth in light conditions and calculated aspect ratios (Fig 4A).

To estimate the change in aspect ratio upon Cdc42 lateral recruitment while taking into account the initial shape of the cell, we normalized the aspect ratios from cells recruiting Cdc42 to those expressing only CRY2 (Fig 4B). CRY2-Cdc42^Q61L,ΔCaaX^ led to a >2-fold reduction in aspect ratio in WT, *rga6Δ*, *rdi1Δ* and *gap1Δ* cells, but had less effect on cell shape change in single and double *rga4Δ* mutants, perhaps in part due to the already wider cell shape of *rga4Δ* cells (Das et al., 2007). Interestingly, CRY2-Cdc42^ΔCaaX^ had little effect on aspect ratio in WT or any single mutants, except in *rga4Δ* cells, which became significantly rounder. Similar, more marked effects were also observed in combinations of *rga4Δ* with *rga6Δ* or *rdi1Δ*. The effect of CRY2-Cdc42^ΔCaaX^ on the shape of these mutants can also readily be observed in plots of cell width to cell length, with the CRY2-Cdc42^ΔCaaX^ cell population placed at an intermediate position between the negative CRY2 and positive CRY2-Cdc42^Q61L,ΔCaaX^ controls (Fig 4C). These data indicate that the optogenetic-dependent Cdc42 activation is counteracted by Rga4 GAP placed at cell sides.

In summary, the data presented in this work show that CRY2-clustered Cdc42 is ectopically activated at cell sides. It is possible that CRY2-tagging interferes with Cdc42 GTPase activity, such that CRY2-Cdc42^ΔCaaX^ represents a slightly activated Cdc42 allele. Alternatively, the CRY2-dependent clustering may lead to Cdc42 activation. This mirrors similar observations made with other GTPases, such as Rac1 and RhoA in mammalian cells (Bugaj et al., 2013). CRY2-mediated Cdc42 activation is not dependent on either individual Cdc42 GEF, though it is possible that both play a redundant role. The activity level of CRY2-Cdc42 at cell sides is sufficient to recruit the GEF Scd1, whose recruitment to Cdc42-GTP depends on the scaffold Scd2 (Lamas et al., 2020), but positive feedback does not appear to become established as Cdc42 activity levels at cell sides are not altered in absence of Scd2. It is possible that the non-physiological linkage of Cdc42 to the plasma membrane (through RitC-CIBN-CRY2 binding instead of through the normal prenyl group) undermines the positive feedback, although we previously showed that a similarly engineered constitutively active Cdc42 allele did trigger feedback-dependent recruitment of endogenous Cdc42 (Lamas et al., 2020). Alternatively, cellular regions may not be equally permissive to Cdc42 activity feedback amplification. For instance, the absence of Ras1 activity on cell sides may reduce the effectiveness of the feedback (Lamas et al., 2020). Conversely, we have shown here that the Cdc42 GAP Rga4, which is enriched on cell sides, antagonizes the effect of CRY2-Cdc42 on cell morphogenesis. Additional mechanisms preventing growth on cell sides are also being proposed (Miller et al, BioRxiv). Thus, one critical question for future research is to better understand the many layers that confer robustness to cell morphogenesis.

## Materials and methods

### Strains, Media, and Growth Conditions

Strains used in this study are listed in Table S1. Standard genetic manipulation methods for *S. pombe* transformation and tetrad dissection were used to generate the strains listed. For microscopy experiments, cells were first pre-cultured in 3 mL of Edinburgh minimal media (EMM) in dark conditions at 30°C for 6 – 8 h. Once exponentially growing, pre-cultures were diluted (Optical Density (O.D.) _600nm_ = 0.02) in 10 mL of EMM and incubated in dark conditions overnight at 30°C. In order to allow proper aeration of the culture, 50 mL Erlenmeyer flasks were used. For cell size analyses cells were pre-cultured and diluted once in 3 mL of Edinburgh minimal media (EMM) in dark conditions at 30°C for 6 – 8 h. Once exponentially growing, pre-cultures were diluted in 10 mL of EMM and incubated in light conditions for a minimum of 14 h. All live-cell imaging was performed on EMM-ALU agarose pads, except calcofluor-white experiments in which cells were placed directly on a slide (Dudin et al., 2015). Gene tagging was performed at endogenous genomic locus at the 3’ end, yielding C-terminally tagged proteins, as described (Bähler et al., 1998). Pak1 gene tagging was performed by transforming a WT strain with AfeI linearized (pBSII(KS^+^))-based single integration vector (pAV72-3’UTR^*pak1*^-Pak1-sfGFP-kanMX-5’UTR^*pak1*^) targeting the endogenous locus. The functional mCherry-tagged and sfGFP-tagged Cdc42 alleles Cdc42-mCherry^sw^ and Cdc42-sfGFP^sw^ were used as described in (Bendezu et al., 2015). Gene tagging, deletion and plasmid integration were confirmed by diagnostic PCR for both sides of the gene.

Construction of plasmids and strains expressing CIBN-mTagBFP2-Ritc, CRY2, CRY2-Cdc42^ΔCaaX^ and CRY2-Cdc42^Q61L,ΔCaaX^ was done as described in (Lamas *et al.* 2020).

To generate the *Pact1-cdc42-mCh*^*SW*^ strain a pINT-ura4^+^ integrative vector was generated. *Pact1* was amplified from gDNA using primers osm2378 (atg<u>ggccc</u>gctagcatgcGATCTACGATAATGAGACGG) and osm2379 (ccgg<u>ctcgag</u>GGTCTTGTCTTTTGAGGGTT) and cloned using ApaI and XhoI. *Cdc42-mCh^SW^-nmt_terminator_* was amplified from pSM1224 using osm2343 (ccc<u>aagctt</u>ATGCCCACCATTAAGTGTGTCG) and osm2344 (gc<u>tctaga</u>CTTCTAATTACACAAATTCCG) and cloned using HindIII and XbaI. As a results pSM1449 was generated. This plasmid was linearized with AvrII and integrated at *ura4* locus of YSM485 strain. The endogenous allele of *cdc42* was deleted using a hygromycin (*hph^+^*) resistance cassette as described (Bähler et al., 1998). *Hph*^*+*^ cassette was amplified from pSM693 using osm2511 (TACTTAGGGGTTTGAACTTTCTAGGAATTCAATAAAGTGAAGCAAAGCTTTACGATTAATTATTTTTTGTGAAATAGTcggatccccgggttaattaa) and osm2512 (AAGCTAAGACATTGTTTACTGTTGTAAACTAGCTGTATTAAGGAAATTTCGGAAAAGGAAAGAAAACCAGGGGTTAAAgaattcgagctcgtttaaac). Finally, strain YSM3732 was generated by transforming the *Cdc42-mCh^SW^-nmt_terminator_* strain with the *hph^+^* resistance cassette.

In primer sequences, restriction sites are underlined. Plasmid maps are available upon request.

### Cell length and width measurements

For cell length and width measurements shown in Fig 4 and Fig S1, cells were grown at 30°C in 10 ml EMM in light and dark conditions respectively. Exponentially growing cells were stained with calcofluor to visualize the cell wall and imaged on a Leica epifluorescence microscope with 60x magnification platform described previously (Dudin et al., 2015). Measurements were made with ImageJ on septating cells. For each experiment strains with identical auxotrophies were used.

Aspect ratios were calculated as:

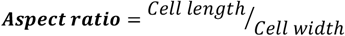

The index of ratios shown in Fig 4B was calculated as:

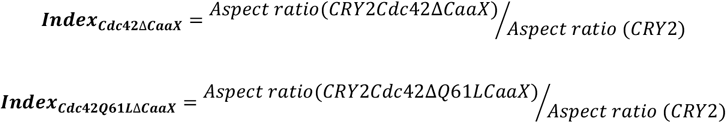

### Microscopy

Fluorescence microscopy experiments were done in a spinning disk confocal microscope, essentially as described (Bendezu and Martin, 2011; Dudin et al., 2015). Image acquisition was performed on a Leica DMI6000SD inverted microscope equipped with an HCX PL APO 100X/1.46 numerical aperture oil objective and a PerkinElmer Confocal system. This system uses a Yokagawa CSU22 real-time confocal scanning head, solid-state laser lines and a cooled 14-bit frame transfer EMCCD C9100-50 camera (Hamamatsu) and is run by Volocity (PerkinElmer). When imaging strains expressing the CRY2-Cdc42^ΔCaaX^ and/or CRY2 systems, an additional long-pass color filter (550 nm, Thorlabs Inc, USA) was used for bright-field (BF) image acquisition, in order to avoid photo-activation by white light.

Spinning disk confocal microscopy experiments shown in Fig 1, 2 and 3 were carried out using cell mixtures (Lamas *et al.* 2020). Cell mixtures were composed by one strain of interest (the sample optogenetic strain, expressing or not an additional GFP-tagged protein) and 2 control strains, namely:

1. <u>RFP control:</u> An RFP bleaching correction strain, expressing cytosolic CRY2PHR-mCherry.
2. <u>GFP control:</u> A wild type strain expressing the same GFP-tagged protein as the strain of interest but without the optogenetic system. This strain was used both as negative control for cell side re-localization experiments and as GFP bleaching correction strain (in Fig 1–2–3 and related supplementary figures).

Strains were handled in dark conditions throughout. Red LED light was used in the room in order to manipulate strains and to prepare the agarose pads. Strains were cultured separately. Exponentially growing cells (O.D._600nm_ = 0.4 – 0.6) were mixed with 2:1:1 (strain of interest, RFP control and GFP control) ratio, and harvested by soft centrifugation (2 min at 1,600 rpm). 1 μL of the cell mixture slurry was placed on a 2 % EMM-ALU agarose pad, covered with a #1.5-thick coverslip and sealed with VALAP (vaseline, lanolin and paraffin). Samples were imaged after 5 – 10 minutes of rest in dark conditions.

The plasma membrane recruitment dynamics of CRY2-Cdc42^ΔCaaX^ and CRY2 systems were assessed using cell mixtures. Protein recruitment dynamics were assessed by applying the 3 different photo-activating cycles listed below. Lasers were set to 100 %; shutters were set to maximum speed and in all instances the RFP channel was imaged first, before the GFP channel. The duration of the experiment was equal regardless of the exposure time settings (≈ 15 s):

- <u>50 ms:</u> RFP channel (200 ms), GFP channel (50 ms). This constitutes one cycle (≈ 0.5 s). 30 time points were acquired (≈ 0.5 s * 30 = 15.1 s).
- <u>250 ms:</u> RFP channel (200 ms), GFP channel (250 ms). This constitutes one cycle (≈ 0.7 s). 22 time points were acquired (≈ 0.7 s * 22 = 15.1 s).
- <u>500 ms:</u> RFP channel (200 ms), GFP channel (500 ms). This constitutes one cycle (≈ 0.9 s). 17 time points were acquired (0.9 s * 17 = 15.5 s).

Endogenous GFP-tagged protein re-localization experiments were carried out using cell mixtures. Lasers were set to 100 %; shutters were set to sample protection and in all instances the RFP channel was imaged first and then the GFP channel. RFP exposure time was always set to 200 ms, whereas the GFP exposure time varied depending on the monitored protein. Cells were monitored in these conditions for 90 s.

Spinning disk confocal time (sum) projections of five consecutive single plane images are shown in Fig S2. Z-stack images were acquired on a Spinning disk confocal microscope using an optimal z-spacing of 0.71 μm between successive stacks. 6 stacks were acquired (covered Z distance= 4 μm).

### Image Analysis

All image-processing analyses were performed with Image J software (http://rsb.info.nih.gov/ij/). Image and time-lapse recordings were imported to the software using the Bio-Formats plugin (http://loci.wisc.edu/software/bio-formats). Time-lapse recordings were aligned using the StackReg plugin (http://bigwww.epfl.ch/thevenaz/stackreg/) according to the rigid body method. All optogenetic data analyses were performed using MATLAB (R2019a), with scripts developed in-house.

### CRY2-Cdc42^ΔCaaX^ and CRY2 quantifications

The plasma membrane recruitment dynamics of CRY2-Cdc42^ΔCaaX^ and CRY2 systems was assessed by recording the fluorescence intensity over a 15 pixel long by 36 pixel wide ROI (roughly 1.25 μm by 3 μm), drawn perpendicular to the plasma membrane of sample cells, from outside of the cell towards the cytosol. The fluorescence intensity values across the length of the ROI were recorded over time in the RFP channel, in which each pixel represents the average of the width (36 pixels) of the ROI (3 replicates, 30 cells per replicate). Average background signal was measured from tag-free wild-type cells incorporated into the cell mixture. The total fluorescence of the Control RFP strain was also measured over time in order to correct for mCherry fluorophore bleaching. In both cases, the ROI encompassed whole cells, where ROI boundaries coincide with the plasma membrane.

Photobleaching correction coefficient was calculated by the following formula:

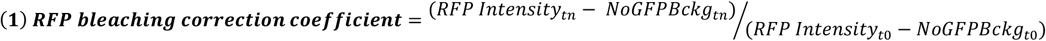

where *RFP intensity* is the signal measured from single RFP control cells, *NoGFPBckg* is the average background signal measured from tag-free cells, t_n_ represents a given time point along the time course of the experiment and t_0_ represents the initial time point (n = 30 time points). These coefficients were corrected by a moving average smoothing method (moving averaged values = 5). RFP bleaching correction coefficient values calculated for individual RFP control cells were averaged in order to correct for bleaching of the RFP signal.

The fluorescence intensity values of optogenetic cells were corrected at each time point with the following formula:

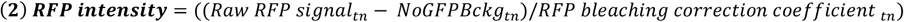

where *Raw RFP signal* is for the RFP values measured from sample strains, *NoGFPBckg* is the average background signal measured from tag-free cells and t_n_ represents a given time point along the time course of the experiment (n = 30 time points). The profiles resulting from these analyses were used to get the net plasma membrane recruitment profiles (Fig S1A, Lamas *et al.* 2020), the fluorescence intensities from the peak ± 1 pixel were averaged and plotted over time.

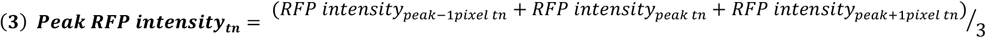

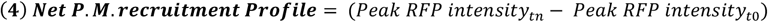

Finally, the single-cell plasma membrane recruitment half-times were calculated by fitting the normalized recruitment profiles with the following formula:

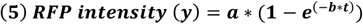

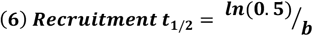

### Quantifications of the re-localization of GFP-tagged proteins to cell sides

Endogenous GFP-tagged protein re-localization was assessed upon photo-activation of CRY2-Cdc42^ΔCaaX^ and CRY2 systems by recording the fluorescence intensity over a 3 pixel-wide by 36 pixel-long (≈ 0.25 μm by 3 μm) ROI drawn parallel to the cell side cortex of sample cells. The average fluorescence intensity values of both GFP and RFP channels were recorded over time from sample strains. In these particular experiments, a GFP control strain was included. These strains serve 2 purposes:

- Calculation of the GFP bleaching correction coefficient (see below).
- <u>Negative control of the experiment.</u> These strains carry the same endogenous GFP-tagged protein as the sample strain of the experiment, however lacking the optogenetic system. This controlled that GFP fluorescence changes were due to the optogenetic system and not caused by imaging per se. Control GFP strains were imaged in the same pad and analyzed in the same way as optogenetic cells.

To derive photobleaching correction coefficients, the average camera background signals (*Bckg*) from 5 cell-free regions was measured as above, and fluorophore bleaching from RFP control and GFP control strains were measured at the cell side of control RFP and control GFP strains, for RFP and GFP channels, respectively.

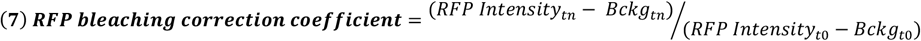

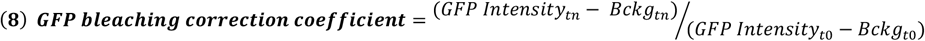

where *RFP intensity* and *GFP intensity* stand for the signal measured from RFP control and GFP control cells, respectively, t_n_ represents a given time point along the time course of the experiment and t_0_ represents the initial time point (n = 30 time points). These coefficients were corrected by a moving average smoothing method, as above.

The fluorescence intensity values of optogenetic cells in both GFP and RFP channels were independently analyzed as follows. First, GFP and RFP signals were background and bleaching corrected, using formulas (7) and (8) for the RFP and GFP channels, respectively:

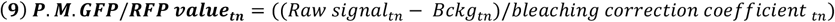

where *Raw signal* intensity represents the GFP or RFP raw values at the cell side cortex, *Bckg* stands for the average fluorescence intensity of 5 independent cell-free regions and t_n_ represents a given time point along the time course of the experiment (n = 30 time points). The net fluorescence intensity at the cell side cortex was then calculated for both GFP and RFP signals.

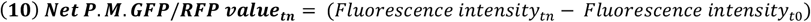

From here on, RFP and GFP signals were treated differently. Single cell plasma membrane RFP profiles from equation (10) were individually normalized and fitted to the equation (5) in order to extrapolate the parameter b. Using the equation (6), recruitment half times of CRY2 and CRY2-Cdc42^ΔCaaX^ were calculated. Because of lower signal-to-noise of the weak GFP fluorescence, plasma membrane GFP profiles from equation (10) were averaged (n > 20 profiles per experiment), and the initial 45 s of the average profile used to extract the half-time of plasma membrane re-localization of endogenous GFP-tagged proteins using equations (5) and (6). 3 experimental replicates were performed and are plotted on Fig 1B; Fig 2B-D, Fig 3B-E-G.

### Quantifications of the re-localization of GFP-tagged proteins from cell tips

Scd1-3GFP tip signal analyses (Fig 3C-F) were performed from the same time-lapse recordings as cell side re-localization experiments. Scd1-3GFP tip signal was recorded over a 3 pixel-wide by 6-12 pixel-long (≈ 0.25 μm by 0.5-1 μm) ROI drawn at the tip of the cells. To derive photobleaching correction coefficients, the average camera background signals (*Bckg*) from 5 cell-free regions was measured as before, and GFP bleaching from GFP control strain was measured at the cell tip.

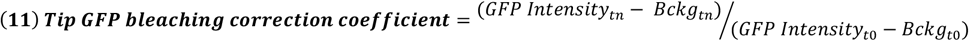

where *GFP intensity* stands for the signal measured from the tip of GFP control cells, t_n_ represents a given time point along the time course of the experiment and t_0_ represents the initial time point (n = 30 time points). This coefficient was corrected by a moving average smoothing method, as before.

The tip GFP fluorescence intensity values of optogenetic cells was analyzed as follows. First, GFP signals was background and bleaching corrected, using formula (12):

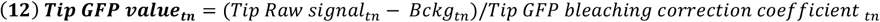

where *Tip Raw signal* intensity represents the GFP raw values at the cell tip, *Bckg* stands for the average fluorescence intensity of 5 independent cell-free regions and t_n_ represents a given time point along the time course of the experiment (n = 30 time points). The tip fluorescence intensities of single optogenetic strains were then normalized relative to their GFP values at the initial time-point.

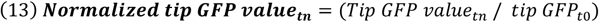

Eventually, average Scd1-3GFP tip signal was calculated (>15 cells, Fig 3C-F).

### Quantifications of CRIB-3GFP and Cdc42-mCh^SW^ relative expression and distribution profiles

CRIB-3GFP fluorescence intensity was measured from sum projection of 6 Z-stacks (Fig S2B). The background signal from cell-free regions were used to correct the data. The relative fluorescence intensities were calculated by dividing the single-cell CRIB-3GFP fluorescence intensity measurements of WT and *Pact1-cdc42-mCh*^*SW*^ cells by the average CRIB-3GFP fluorescence intensity of WT cells.

CRIB-3GFP distribution profiles were generated from sum projection images of 5 middle-sections (Fig S2C). 3-pixel wide ROIs were drawn from side to side following cell membrane contour. The background signal from cell-free regions were used to correct the data. Whole tip profiles were split in half based on the pixel position of their maximum CRIB-3GFP intensity (approximately in the middle of the profile), generating 60 half tips. To take into account the CRIB-3GFP expression level, the average CRIB-3GFP profiles from WT and *Pact1-cdc42-mCh*^*SW*^ cells were then normalized by dividing each value by the relative CRIB-3GFP fluorescence intensities values shown in Figure S2B.

### Cell size measurements, aspect ratios and index of ratios

The aspect ratio of mutant cells was calculated by dividing the cell length by the cell width (Fig 4A). In Fig 4B, the index of ratios were calculated by dividing CRY2 control aspect ratios by CRY2-Cdc42^ΔCaaX^ and CRY2-Cdc42^Q61L,Δ^CaaX.

Figures were assembled with Adobe Photoshop CS5 and Adobe Illustrator CS5. All error bars on bar graphs are standard deviations. For statistical analysis, in Fig 1, 2 and 3, cumulative GFP signal (addition of GFP signal along the 30 time points of the time-lapse) was calculated from single cell traces of CRY2, CRY2-Cdc42^ΔCaaX^ and GFP control cells. For statistical analysis, single cell cumulative GFP signals of the entire dataset (3 independent experiments combined) were considered, without averaging. Data normality was assessed by the Lilliesfors test and significance by pair-wise Kruskal-Wallis analysis. P values show significance of differences between CRY2-Cdc42^ΔCaaX^ and CRY2 cells, unless indicated otherwise. T-test was used in Fig 4. On each box the central red mark indicates the median, while the bottom and the top edges indicate the 25^th^ and 75^th^ percentiles respectively, the whiskers extend to the most extreme data points not considering outliers, which are plotted individually using the red ‘+’ symbol. All experiments were done at least three independent times.

## Acknowledgements

We thank Dr Aleksandar Vjestica (University of Lausanne) for gift of the CRY2-Cdc42^ΔCaaX^ strain and Dr Serge Pelet (University of Lausanne) for help with MatLab. This work was funded by ERC Consolidator grant (CellFusion) to SGM.

## Author contributions

IL and SGM conceived the project. NW performed the experiments presented in Fig 4. IL performed all other experiments. IL and SGM wrote the manuscript. SGM acquired funding.

---

**Figure S1.**
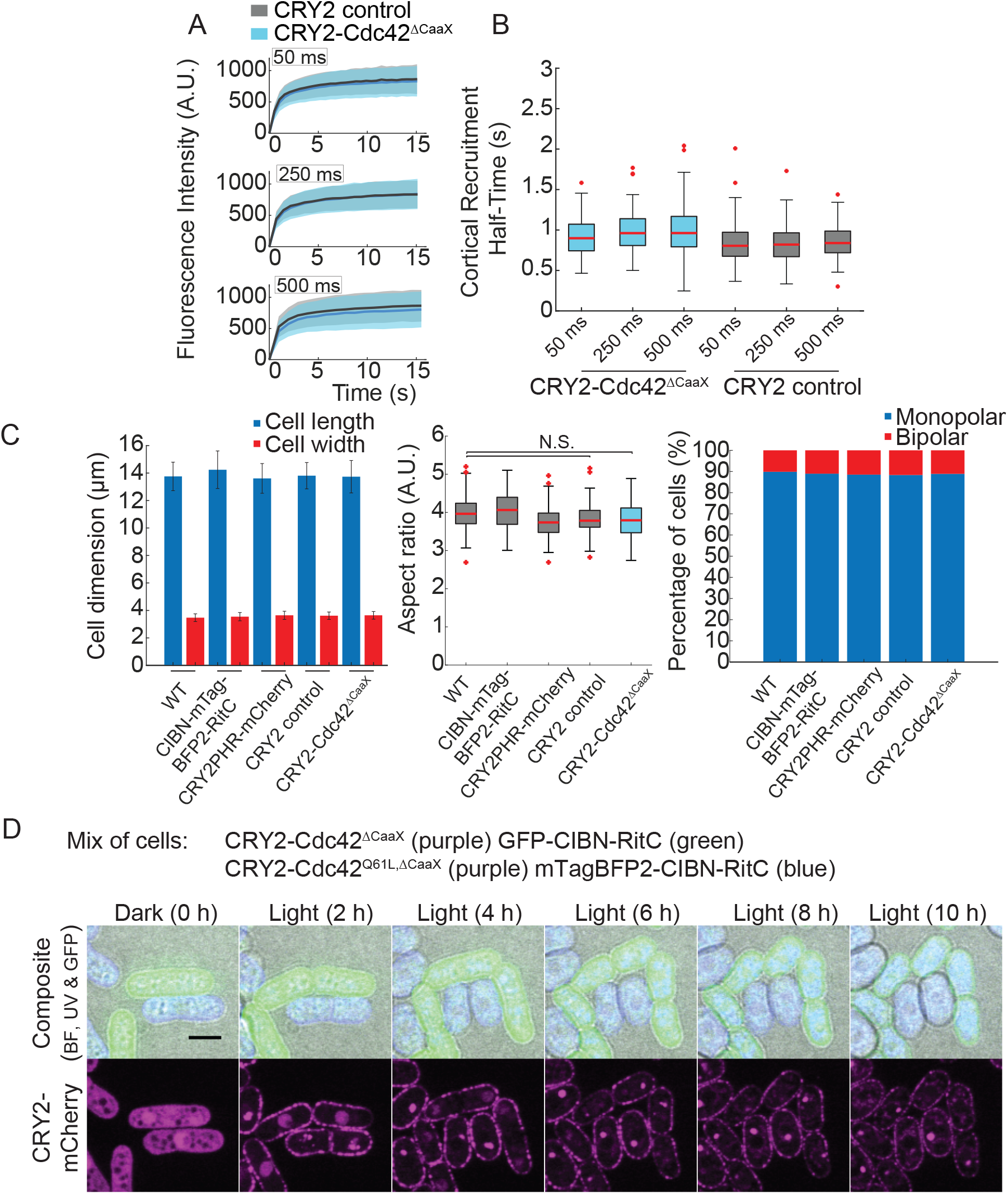
Controls for CRY2-Cdc42^ΔCaaX^ optogenetic recruitment. (A) Plasma membrane recruitment dynamics of CRY2 and CRY2-Cdc42^ΔCaaX^ in response to periodic 50 ms (top), 250 ms (middle) and 500 ms (bottom) blue-light (λ = 485 nm) pulses (N = 3; n = 30 cells per experiment). Thick line = average; shaded area = standard deviation. CRY2 values are from (Lamas et al., 2020). (B) Plasma-membrane recruitment half-times for CRY2 and CRY2-Cdc42^ΔCaaX^. On each box, the central mark indicates the median; the bottom and the top edges indicate the 25th and 75th percentiles, respectively; the whiskers extend to the most extreme data points not considering outliers, which are plotted individually using the red “+” symbol. (C) Cell length and width measurements, aspect ratio and bipolarity of calcofluor-stained cells growing in the dark. CRY2 and CRY2-Cdc42^ΔCaaX^ do not have any effect on cell morphology (p^WTvsCRY2^ = 0.19; p^WTvsCRY2-Cdc42^ = 0.19; p^CRY2vsCRY2-Cdc42^ = 0.83). Monopolarity and bipolarity were assessed on septating cells. (D) Blue light–dependent induction of isotropic growth in CRY2-Cdc42^Q61LΔCaaX^ (blue cells), but not CRY2-Cdc42^ΔCaaX^ (green cells) cells photoactivated at 10-min interval (GFP, RFP, and BF channels were acquired every 10 min; UV channel every 1 h). Note that the patchy appearance of CRY2 is likely due to the clustering properties of this protein. Bars = 5μm.

**Figure S2.**
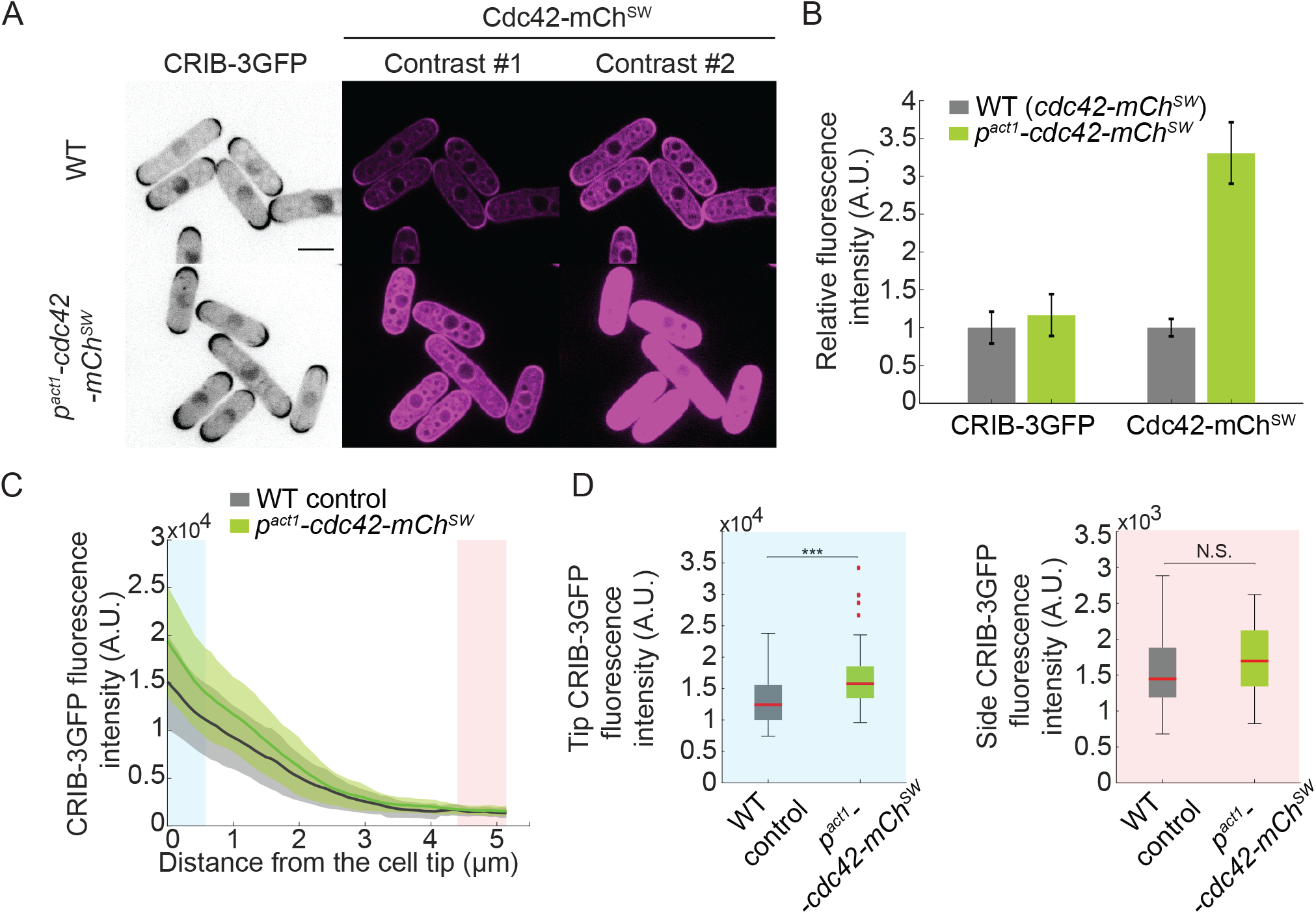
Overexpression of Cdc42-mCh^SW^ in WT cells does not induce ectopic activation of Cdc42. (A) *p*^*act1*^ promoter induces overexpression of Cdc42-mCh^SW^ in CRIB-3GFP cells. Two different contrasting settings are shown. (B) Relative expression of CRIB-3GFP and Cdc42-mCh^SW^ in WT (*cdc42-mCherry^SW^*) and *p*^*act1*^*-cdc42-mCherry*^*SW*^ cells. *p^act1^-cdc42* cells exhibit 1.2x and 3.3x higher CRIB-3GFP and Cdc42-mCh^SW^ fluorescence intensities respectively as compared to WT cells. *N* = 1; *n* = 50 cells. (C) Distribution profile of CRIB-3GFP in WT and *p^act1^-cdc42* cells at the cortex from the centre of cell tips towards the cell side. Blue area represents the centre of the tip (0.5 μm) and red area represents the cell side. The CRIB profile was corrected by the 1.2-fold increase in CRIB expression. CRIB-3GFP fluorescence intensities at the cell tip (left) and cell sides (right) in WT (*cdc42-mCherry^SW^*) and *p*^*act1*^*-cdc42-mCherry*^*SW*^ cells derived from the blue and red areas shown in C. p^CRIB-3GFP-Tip^ = 3.7e-5; p^CRIB-3GFP-Side^ = 0.18. *N* = 1; *n* = 60 half tips. In all graphs, thick line = average; bars & shaded area = standard deviation; WT, wild type; A.U., arbitrary units. Bars = 5 μm.

**Figure S3.**
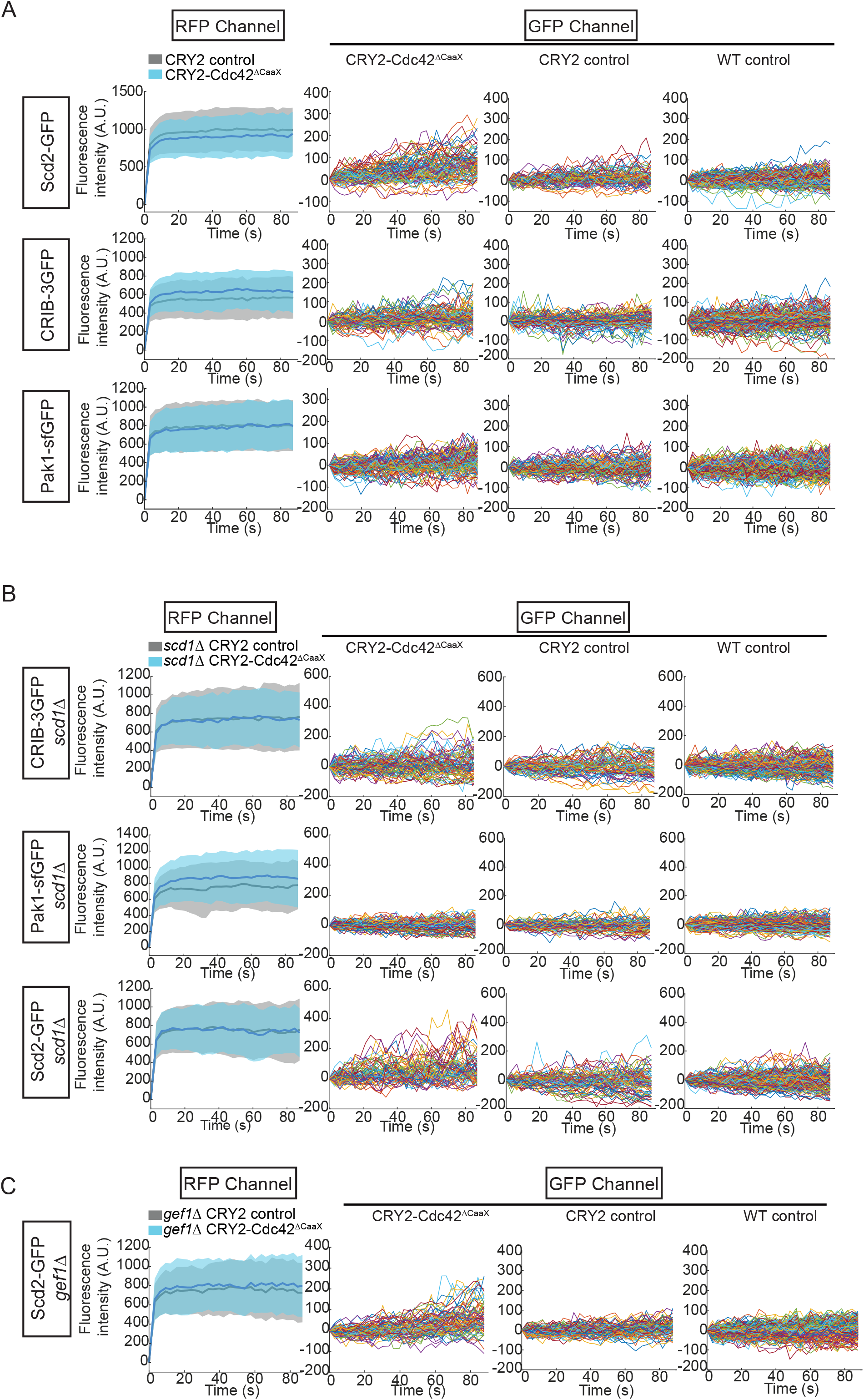
Control and single-cell traces for optogenetic recruitment in WT, *scd1Δ and gef1Δ* cells. (A) Single-cell traces corresponding to the average plots shown in Fig 1B. The left column shows the average RFP signal at the plasma membrane of wildtype CRY2-Cdc42^ΔCaaX^ and CRY2 cells. The three other graphs show, from left to right, single cell GFP traces of CRY2-Cdc42^ΔCaaX^, CRY2 and WT control cells for Scd2-GFP, CRIB-3GFP and Pak1-sfGFP in otherwise wildtype cells. (B) Single-cell traces corresponding to the average plots shown in Fig 3B. The left column shows the average RFP signal at the plasma membrane of wildtype CRY2-Cdc42^ΔCaaX^ and CRY2 cells. The three other graphs show, from left to right, single cell GFP traces of CRY2-Cdc42^ΔCaaX^, CRY2 and WT control cells for CRIB-3GFP, Pak1-sfGFP and Scd2-GFP in *scd1Δ* mutants. (C) Single-cell traces corresponding to the average plots shown in Fig 3D. The left column shows the average RFP signal at the plasma membrane of wildtype CRY2-Cdc42^ΔCaaX^ and CRY2 cells. The three other graphs show, from left to right, single cell GFP traces of CRY2-Cdc42^ΔCaaX^, CRY2 and WT control cells for Scd2-GFP in *gef1Δ* mutants.

**Figure S4.**
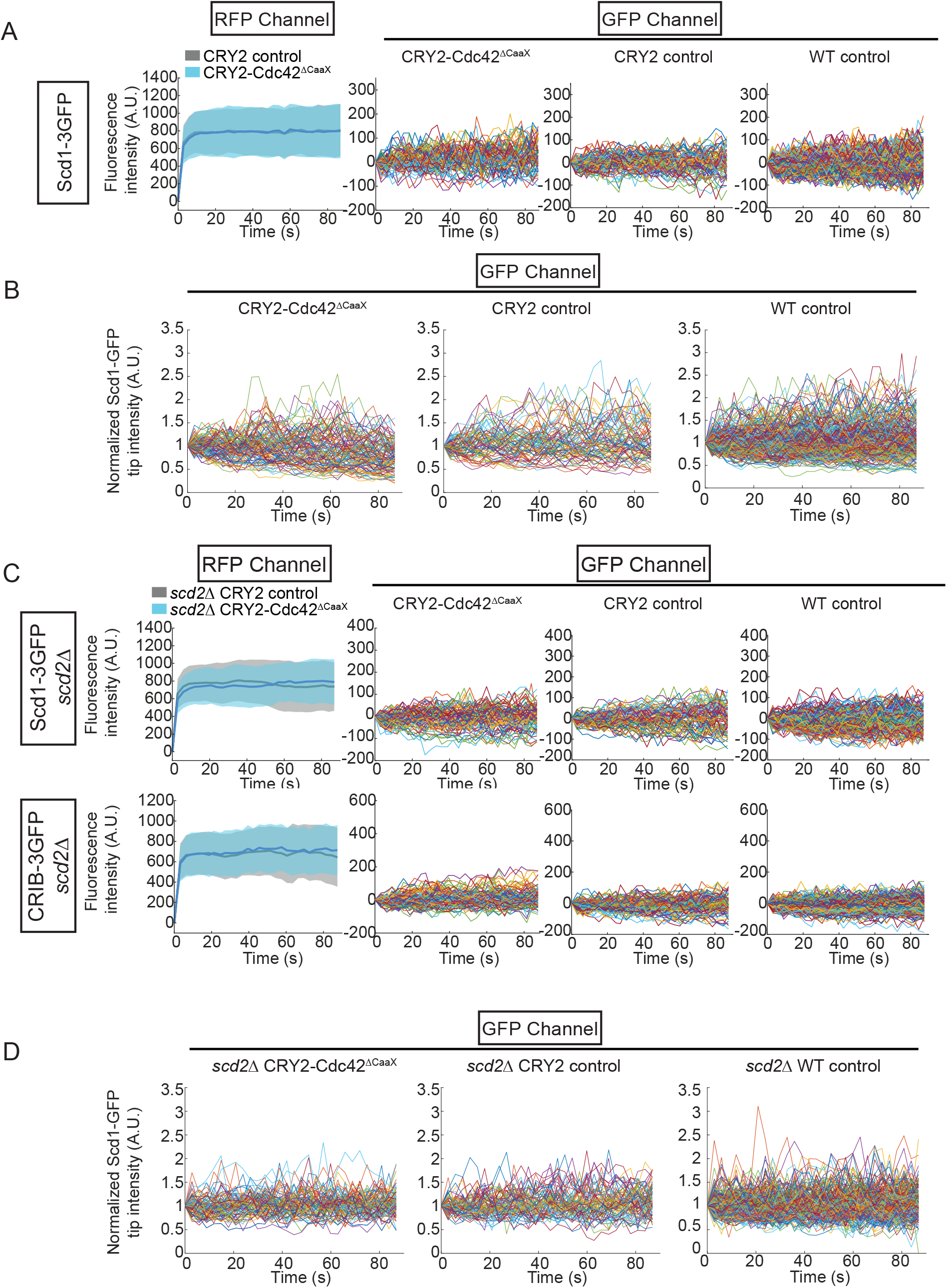
Control and single-cell traces for optogenetic recruitment of Scd1-3GFP and CRIB-3GFP in WT *and sdc2Δ* cells. A) Single-cell traces corresponding to the average plots shown in Fig 4B. The left column shows the average RFP signal at the plasma membrane of wildtype CRY2-Cdc42^ΔCaaX^ and CRY2 cells. The three other graphs show, from left to right, single cell GFP traces of CRY2-Cdc42^ΔCaaX^, CRY2 and WT control cells for Scd1-3GFP in otherwise wildtype cells. (B) Single-cell traces corresponding to the average plots shown in Fig 4C. The three graphs show, from left to right, single cell GFP traces of CRY2-Cdc42^ΔCaaX^, CRY2 and WT control cells for normalized tip signal of Scd1-3GFP in otherwise wildtype cells. (C) Single-cell traces corresponding to the average plots shown in Fig 4E-G. The left column shows the average RFP signal at the plasma membrane of wildtype CRY2-Cdc42^ΔCaaX^ and CRY2 cells. The three other graphs show, from left to right, single cell GFP traces of CRY2-Cdc42^ΔCaaX^, CRY2 and WT control cells for Scd1-3GFP and CRIB-3GFP in *scd2Δ* mutants. (D) Single-cell traces corresponding to the average plots shown in Fig 4F. The three graphs show, from left to right, single cell GFP traces of CRY2-Cdc42^ΔCaaX^, CRY2 and WT control cells for normalized tip signal of Scd1-3GFP in *scd2Δ* mutants.

**Table S1:**
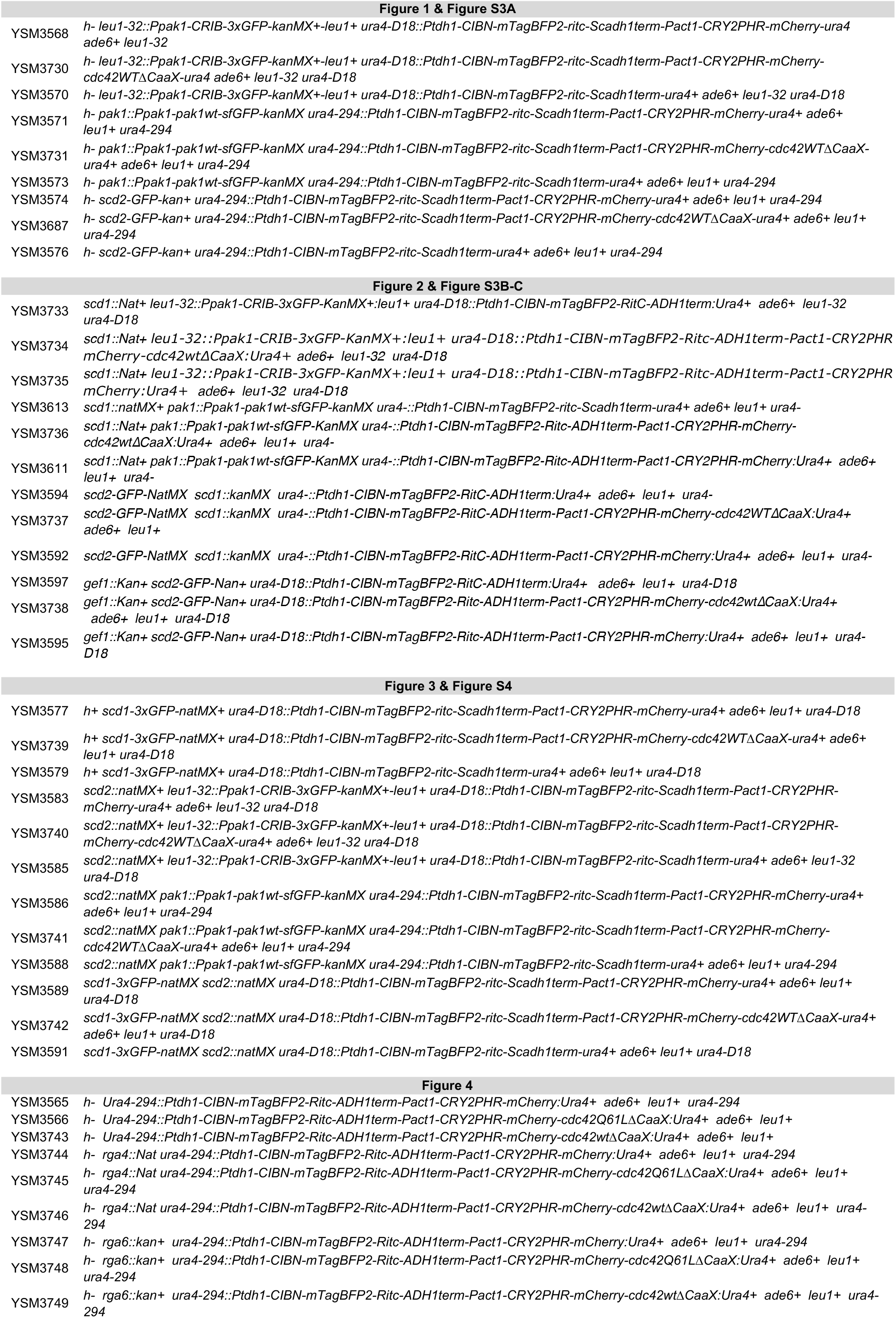
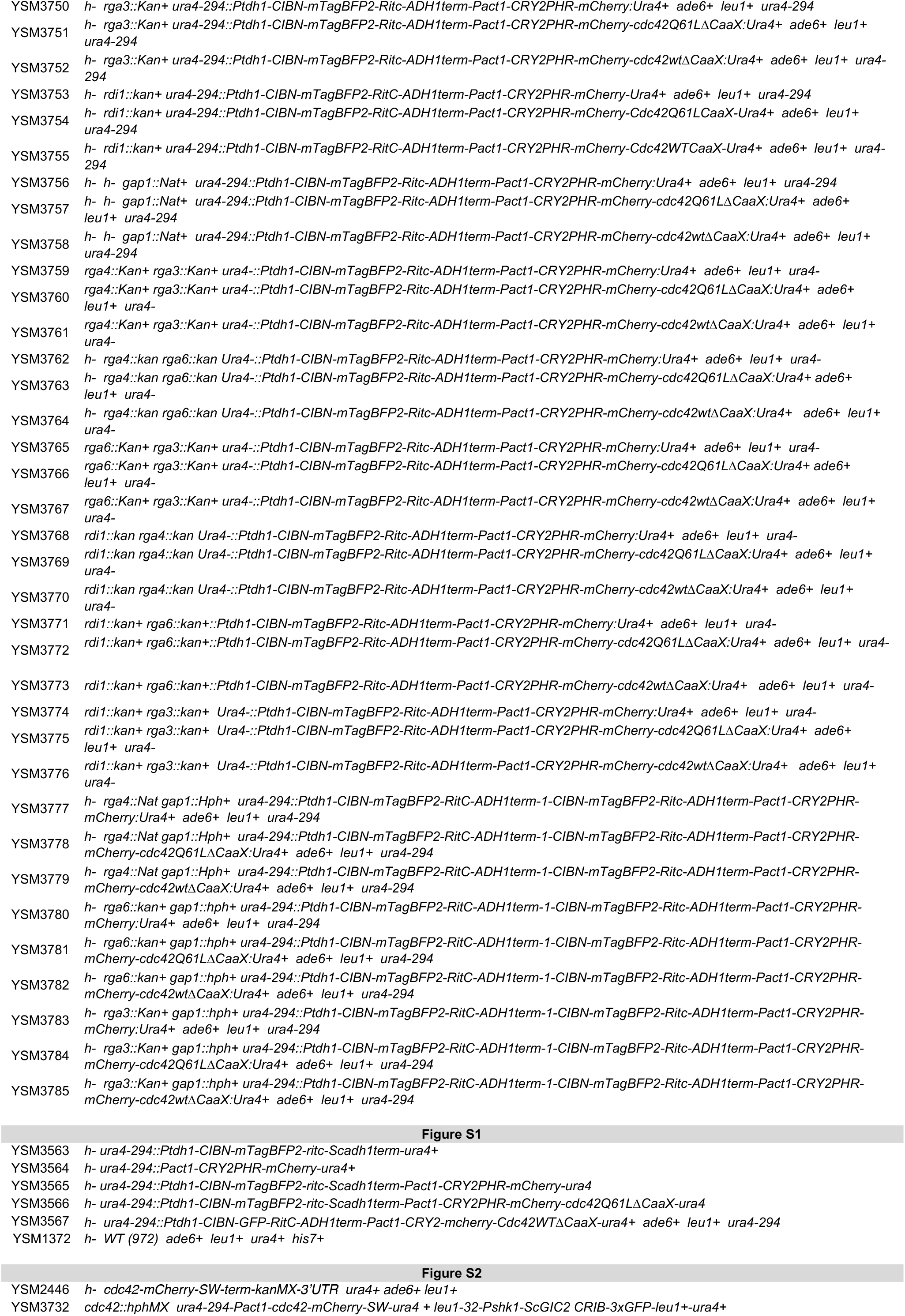
List of strains used in this study.

